# Common genetic variants with fetal effects on birth weight are enriched for proximity to genes implicated in rare developmental disorders

**DOI:** 10.1101/2020.07.02.184028

**Authors:** Robin N. Beaumont, Isabelle K. Mayne, Rachel M. Freathy, Caroline F. Wright

## Abstract

Birth weight is an important factor in newborn and infant survival, and both low and high birth weights are associated with adverse later life health outcomes. Genome-wide association studies (GWAS) have identified 190 loci associated with either maternal or fetal effects on birth weight. Knowledge of the underlying causal genes and pathways is crucial to understand how these loci influence birth weight, and the links between infant and adult morbidity. Numerous monogenic developmental syndromes are associated with birth weights at the extreme upper or lower ends of the normal distribution, and genes implicated in those syndromes may provide valuable information to help prioritise candidate genes at GWAS loci. We examined the proximity of genes implicated in developmental disorders to birth weight GWAS loci at which a fetal effect is either likely or cannot be ruled out. We used simulations to test whether those genes fall disproportionately close to the GWAS loci. We found that birth weight GWAS single nucleotide polymorphisms (SNPs) fall closer to such genes than expected by chance. This is the case both when the developmental disorder gene is the nearest gene to the birth weight SNP and also when examining all genes within 258kb of the SNP. This enrichment was driven by genes that cause monogenic developmental disorders with dominant modes of inheritance. We found several examples of SNPs located in the intron of one gene that mark plausible effects via different nearby genes implicated in monogenic short stature, highlighting the closest gene to the SNP not necessarily being the functionally relevant gene. This is the first application of this approach to birth weight loci, which has helped identify GWAS loci likely to have direct fetal effects on birth weight which could not previously be classified as fetal or maternal due to insufficient statistical power.

## Introduction

Weight at birth is an important factor in newborn and infant survival [1], and is associated with a higher risk of adverse adult health outcomes at both the high and low ends of the population distribution [2–4]. Variation in birth weight is influenced by a combination of environmental and genetic factors, and genome-wide association studies (GWAS) of birth weight have implicated 190 genomic loci to date [5–7]. The associated variants at three-quarters of the identified loci where classification is possible show direct effects of the fetal genotype, while the rest represent SNPs having only indirect effects of the maternal genotype (acting via the intrauterine environment) [5]. Knowledge of the causal genes and biological pathways underlying birth weight variation will be crucial to understanding its links with infant and adult morbidity. However, causal variants at the identified GWAS loci have not yet been identified; many of the single nucleotide polymorphisms (SNPs) that mark the association signals fall outside coding regions and it is unclear whether the functional variant they are tagging exerts its effect via the nearest gene or elsewhere.

Rare developmental syndromes arising from severe mutations in a single, known gene may provide valuable information to help prioritise candidate genes at GWAS loci [8–10]. Numerous monogenic developmental syndromes include either extreme fetal overgrowth (e.g. Cantu syndrome caused by mutations in *ABCC9* [11] and Clove syndrome caused by mutations in *PIK3CA* [12]) or severe fetal growth restriction (e.g. Floating-Harbour syndrome caused by mutations in *SRCAP* [13,14] and Myhre syndrome caused by mutations in *SMAD4* [15,16]). The overlap between genes with monogenic effects on birth weight and loci associated with birth weight from GWAS has not been formally examined.

Following a previous GWAS of adult height, Wood et al. [17] used a curated list of genes associated with rare human conditions of abnormal skeletal growth to investigate the identified loci. They hypothesised that common variation in or near the genes on the list would underlie several of the GWAS signals, and thereby implicate biological pathways of relevance to normal variation in adult height. They found that the height GWAS loci were 1.4-fold more likely to fall near to the curated list of genes than simulated lists of randomly selected SNPs/indels. It is not known whether a similar relationship exists between monogenic and polygenic loci for fetal genetic variation underlying birth weight variation. If such an overlap exists, it could help to prioritise candidate genes at these loci, and to understand the biological pathways underlying birth weight. We tested whether genes known to cause severe developmental disorders [18] were nearer lead birth weight GWAS SNPs with evidence of fetal effects, than expected by chance (Figure 1). We found evidence that the birth weight GWAS SNPs tested fell disproportionately close to genes that cause severe developmental disorders, and that this was driven by disease genes that act via a dominant mechanism. This approach helps to highlight potentially causal genes at GWAS loci, underscored by the fact that, for 24 of the 37 SNPs falling near to developmental disorder genes, the nearest gene to the SNP was not the developmental disorder gene.

**Figure 1.**
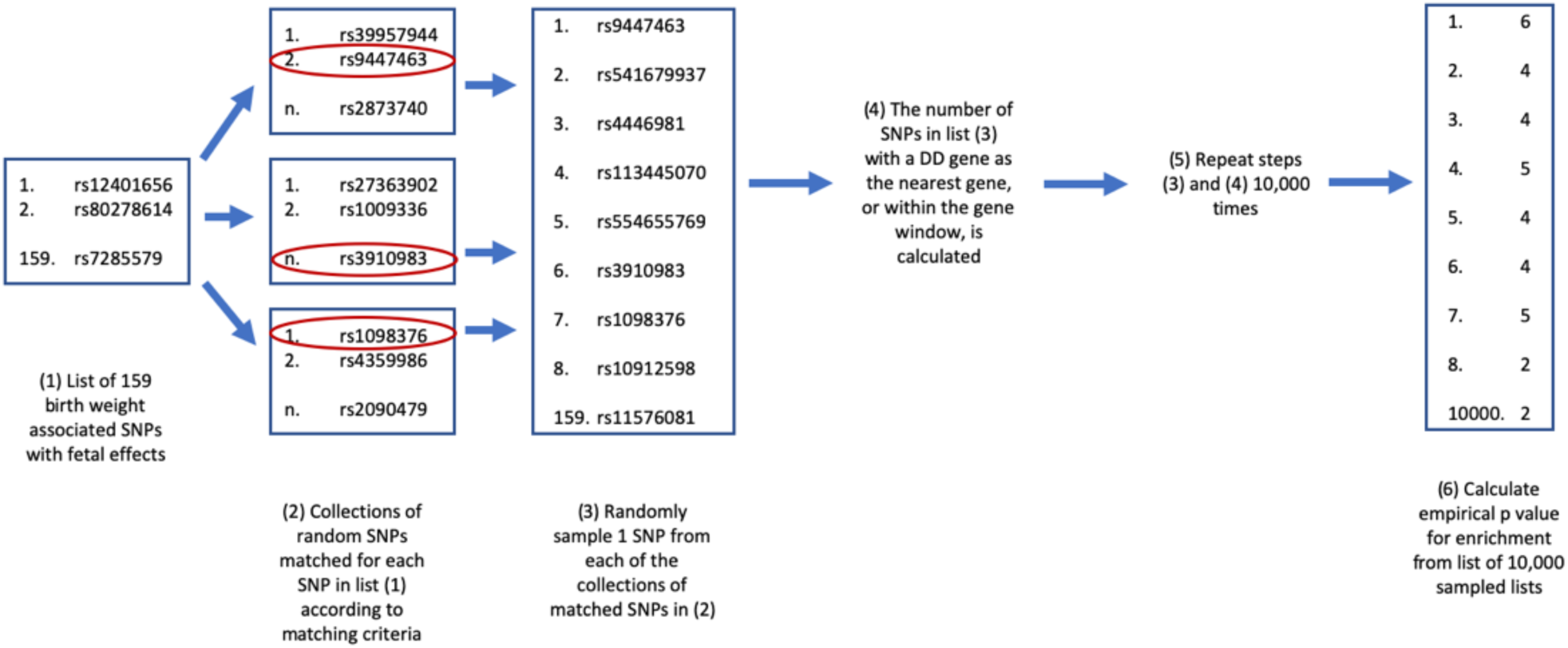
Flow chart showing each of the steps in the enrichment analysis

## Methods

### Birth Weight SNPs

We selected the lead SNP at each of the 190 genomic loci from the latest GWAS of birth weight [5]. Where a locus is known to have different lead SNPs from the maternal GWAS of offspring birth weight versus the GWAS of own birth weight (“fetal GWAS”), we selected the lead SNP from the fetal GWAS. In that study, the 190 loci had been classified into categories according to the likely origin of their effects on birth weight (Table S1): “Fetal only” (62 SNPs); “Maternal only” (31 SNPs); “Fetal and Maternal” (35 SNPs); and “Unclassified” (62 SNPs). Since we were interested in investigating loci with direct fetal effects on birth weight, we excluded the loci classified as “Maternal only” from our analyses. Loci on chromosome X (N=4) were also excluded from our analyses due to the difficulty in classifying X chromosome genes as dominant or recessive. The resulting list of lead SNPs used in our analysis included 159 SNPs.

### Gene Lists

A list of genes definitively linked to monogenic developmental disorders [19] was downloaded from https://www.ebi.ac.uk/gene2phenotype/ on 18 July 2018. Genes on the X-chromosome were excluded. Genes were separated into groups based on the mode of inheritance of their associated developmental disorders (dominant, recessive or both). The list of developmental disorder genes can be found in Table S2.

### Enrichment Analysis

We aimed to test whether our 159 selected lead SNPs, marking common fetal variant effects on birth weight, fall near to genes in which rare variants cause developmental disorders (that may include high or low birth weight) more often than would be expected by chance, i.e. we tested for “enrichment” of proximity to developmental syndrome genes in our list of GWAS SNPs. For each enrichment analysis, we used the 17,073,342 SNPs with minor allele frequency (MAF) >=0.1% included in the UK Biobank HRC-imputed dataset (release v3 March 2018) as a reference [20]. From this list of reference SNPs, we selected 10,000 lists of SNPs which were matched to the lead SNPs from GWAS of birth weight based on the matching criteria listed below. These lists of matching SNPs were used to create an empirical distribution, described below, from which we calculated empirical P values for the corresponding list of birth weight loci (Figure 1). We used two sets of matching criteria: (1) the distance to the nearest gene, and (2) the number of genes within a given distance. We repeated these analyses splitting the list of developmental disorder genes into those with dominant modes of inheritance and those with recessive modes of inheritance. These criteria are described in more detail in the following sections, and the code required to run the analysis has been packaged and can be downloaded from https://github.com/rnbeaumont/DD_gene_enrichment.

### (1) Nearest Gene

Each of the 159 birth weight lead SNPs was annotated with its nearest gene and the distance to that gene. The criteria for selecting 10,000 lists of matched SNPs for the nearest gene analysis were: MAF for the matching SNP between 0.9-1.1x the MAF of the index SNP; and distance to the nearest gene of the matching SNP within ±10% of the distance of the index SNP to the nearest gene. For each of the 10,000 lists of matched SNPs, we calculated the number of SNPs for which their nearest gene appeared in the lists of developmental disorder genes. We also calculated the number of developmental disorder genes that appeared in the nearest gene list for the matched SNPs. These were used as our empirical distributions. We then calculated the number of nearest genes for the birth weight loci which appeared in the developmental disorder genes lists and vice versa.

### (2) Gene Windows

We annotated each birth weight SNP with the number of genes 19kb, 94kb, 138kb and 256kb either side of the SNP. These windows were chosen as the mean, median, lower quartile and upper quartile of the distances from lead birth weight SNPs to eight placenta eQTL genes [21] from the Warrington et al GWAS of birth weight [5] as these represent biologically plausible distances between functional units. The criteria for selecting the 10,000 lists of matched SNPs for the gene window analyses were: the MAF of the matching SNP within 0.9-1.1x the MAF of the lead birth weight SNP; and number of genes within the window matching that of the lead birth weight SNP. For each list of the lists of matching SNPs, we calculated the number of SNPs for which one or more of the genes within the relevant window was in the list of developmental disorder genes, and the number of those genes which appear within the relevant distance of at least one matched SNP. The empirical P values for the number of birth weight SNPs for which at least one of the genes within the window appear in the list of developmental disorder genes and vice versa using the empirical distributions.

### Sensitivity Analysis

These analyses were repeated excluding birth weight SNPs categorised as “Unclassified” [5] as a sensitivity analysis, as that category could include SNPs with maternal effects.

## Results

### Nearest Gene

The full list of genes linked to rare monogenic developmental disorders contained 1362 autosomal genes. Of these, 20 were the closest gene for at least one lead SNP from the GWAS of birth weight. The P value for enrichment compared to the empirical distribution of matching SNPs was P=0.0002 (Table 1). Of the 159 birth weight SNPs, the nearest gene for 22 SNPs was in the full list of developmental disorder genes (P=0.0036) (Table 2). When we split the list of genes into those that cause disease via either a dominant (n=475) or recessive (n=887) mode of inheritance only, 14 dominant genes were the nearest gene of at least one birth weight SNP (P<0.0001) compared to six recessive-only genes (P=0.17). Of the birth weight SNPs, the nearest gene for 15 SNPs (P<0.0001) was in the dominant gene list, and seven (P=0.55) for the recessive-only gene list.

**Table 1:** Number of Developmental Disorder genes which are the nearest gene, or within the corresponding window, of 159 birth weight SNPs annotated as either “Fetal Only”, “Maternal and Fetal”, or “Unclassified”, and the corresponding empirical P value.

|  | Nearest |  | 19kb |  | 94kb |  | 138kb |  | 258kb |  |
| --- | --- | --- | --- | --- | --- | --- | --- | --- | --- | --- |
| Gene List (Number of Genes) | N<sub>genes</sub> | P | N<sub>genes</sub> | P | N<sub>genes</sub> | P | N<sub>genes</sub> | P | N<sub>genes</sub> | P |
| All Genes (1362) | 20 | 2.00E-04 | 22 | 2.10E-03 | 48 | 2.00E-04 | 57 | 1.50E-03 | 82 | 1.00E-03 |
| Dominant (475) | 14 | <1.00E-4 | 13 | <1.00E-4 | 25 | <1.00E-4 | 27 | <1.00E-4 | 36 | 3.00E-04 |
| Recessive (936) | 10 | 1.90E-02 | 13 | 1.90E-02 | 27 | 9.70E-03 | 35 | 7.50E-03 | 51 | 5.90E-03 |
| Recessive excluding Dominant (887) | 6 | 1.68E-01 | 9 | 1.15E-01 | 23 | 2.48E-02 | 30 | 1.67E-02 | 46 | 1.02E-02 |

**Table 2.** Number of birth weight SNPs (total NSNPs = 159) for which a developmental disorder gene is the nearest gene, or for which a developmental disorder gene is within the corresponding window, and the corresponding empirical P value. Birth weight SNPs included are all those classified as classified as “Fetal Only”, “Maternal and Fetal”, or “Unclassified” in the GWAS of birth weight [5].

|  | Nearest |  | 19kb |  | 94kb |  | 138kb |  | 258kb |  |
| --- | --- | --- | --- | --- | --- | --- | --- | --- | --- | --- |
| Gene List (Number of Genes) | N<sub>SNPs</sub> | P | N<sub>SNPs</sub> | P | N<sub>SNPs</sub> | P | N<sub>SNPs</sub> | P | N<sub>SNPs</sub> | P |
| All Genes (1362) | 22 | 3.60E-03 | 24 | 5.00E-03 | 45 | 1.20E-03 | 50 | 1.09E-02 | 67 | 1.12E-02 |
| Dominant (475) | 15 | <1.00E-04 | 14 | 8.00E-04 | 26 | <1.00E-04 | 27 | 1.30E-03 | 37 | 1.30E-03 |
| Recessive (936) | 12 | 8.37E-02 | 15 | 5.53E-02 | 29 | 6.65E-02 | 25 | 7.91E-02 | 48 | 4.42E-02 |
| Recessive excluding Dominant (887) | 7 | 5.47E-01 | 10 | 4.18E-01 | 24 | 2.49E-01 | 29 | 3.35E-01 | 42 | 1.85E-01 |

### Gene Windows

Of the full list of developmental disorder genes, 22, 48, 57 and 82 genes fell within 19kb, 94kb, 138kb and 258kb of at least one birth weight SNP respectively (P=0.0021; P=0.0002; P=0.0015; P=0.001 respectively). Of the birth weight SNPs, 24, 45, 50 and 67 SNPs had at least one gene from the full gene list within 19kb, 94kb, 138kb and 258kb respectively (P=0.006; P=0.0012; P=0.011; P=0.011). Genes in which rare mutations cause dominant developmental disorders showed strong evidence of enrichment within the gene-windows analysis with 13, 25, 27 and 36 genes respectively in the 19kb, 94kb, 138kb and 258kb windows (P<0.0001; P<0.0001; P=0.0001; P=0.0003). Of the birth weight SNPs, 14, 26, 27 and 37 had at least one dominant disease gene within each of the windows (P=0.0008; P<0.0001; P=0.0013; P=0.0013). There was little evidence that genes in which rare mutations cause only recessive disease showed any enrichment for falling with gene windows with 9, 23, 30 and 46 (P=0.12; P=0.025; P=0.010; P=0.017) genes falling within each window, and 10, 24, 29 and 42 (P=0.42; P=0.25; P=0.33; P=0.19) SNPs with at least one recessive-only gene within each window respectively.

Results from sensitivity analysis excluding “Unclassified” birth weight SNPs showed similar patterns (Tables S3 and S4).

## Discussion

This is the first study to investigate the overlap between birth weight GWAS signals and genes known to cause rare developmental disorders. We found that common lead SNPs from GWAS that are associated with birth weight, either partly or entirely through direct fetal effects, fall disproportionately closer to such genes than to randomly-selected similar genes. This enrichment for associations was driven by developmental disorder genes with dominant modes of inheritance, and the pattern was seen both for the nearest gene analysis, and for all window sizes in the gene window analyses.

The interpretation of GWAS loci and the genes and pathways impacted by them for complex traits such as birth weight is less straight forward than that of molecular phenotypes such as urate levels [22]. Rare monogenic variants that cause severe disease are unlikely to underlie the associations with common SNPs that are identified in GWAS [23]. Rather, the lead SNPs are far more likely to tag functional variants of a similar frequency. Genes that are causally linked with any phenotype may harbour a spectrum of genetic variants, from rare with severe consequences (such as complete loss of gene function), to common with mild consequences (such as reduced gene expression). Our results support this hypothesis and show that the genes implicated in rare developmental syndromes can help to prioritise candidate causal genes at birth weight loci. Furthermore, the fact that enrichment patterns were observed for developmental disorders with a dominant mode of inheritance suggests that the mechanism of action of common birth weight variants in the same genes is also likely dominant or additive, rather than recessive (see Table 3).

**Table 3:**
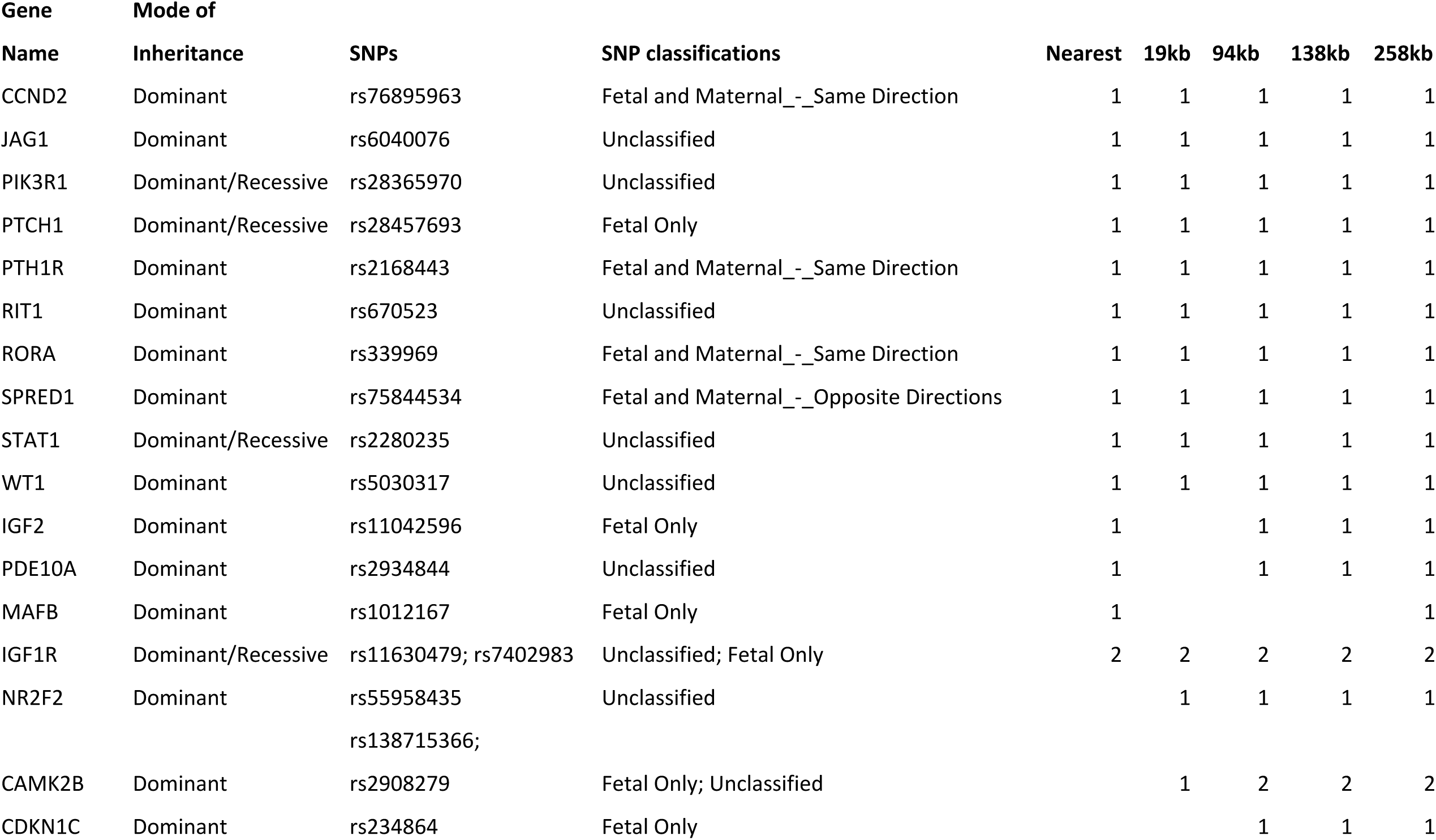
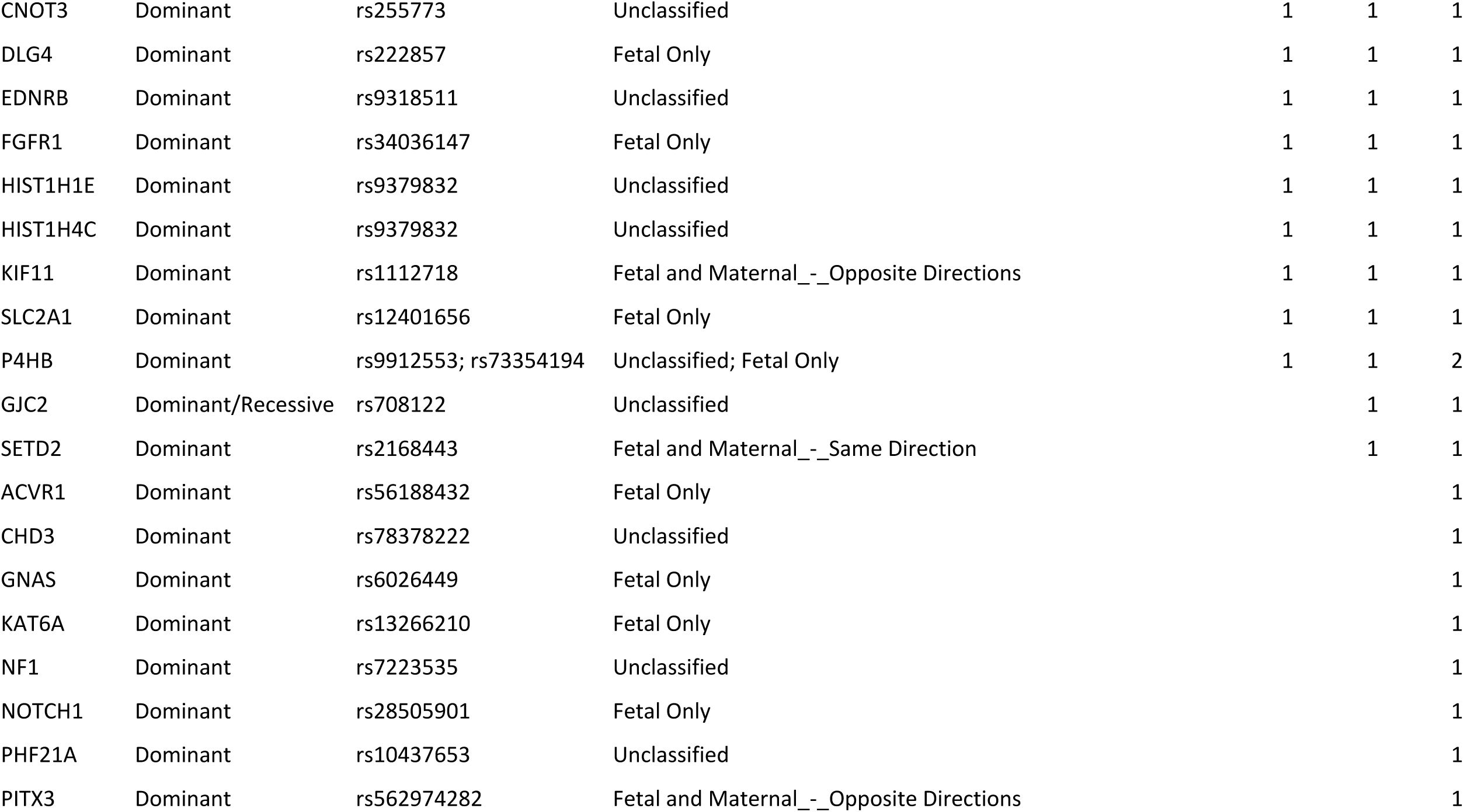
Dominant developmental disorder genes falling near to birth weight SNPs in each of our nearest gene or gene window analyses. Columns 5 to 9 indicate the number of birth weight SNPs for which that gene was the nearest gene or was within the relevant gene window.

Of the 37 birth weight SNPs with developmental disorder genes within the largest 258kb window, this gene is the nearest one for just 13. A histogram of the distance from these SNPs to the developmental disorder gene is shown in Figure 2. While a developmental disorder gene is unlikely to be the relevant functional gene for every birth weight SNP, this result nonetheless highlights the fact that the nearest gene to the SNP is not necessarily the best candidate for functionally relevant genes. Our analysis has also helped categorise GWAS SNPs previously unclassified with respect to maternal or fetal activity and prioritise likely candidate genes. For example, high birth weight is a feature of Noonan syndrome, which can be caused by missense mutations in the *RIT1* gene [24]; one of the birth weight SNPs, “Unclassified” in the recent birth weight GWAS, lies within the gene boundaries of *RIT1*, suggesting that the SNP is acting through the fetal genome.

**Figure 2:**
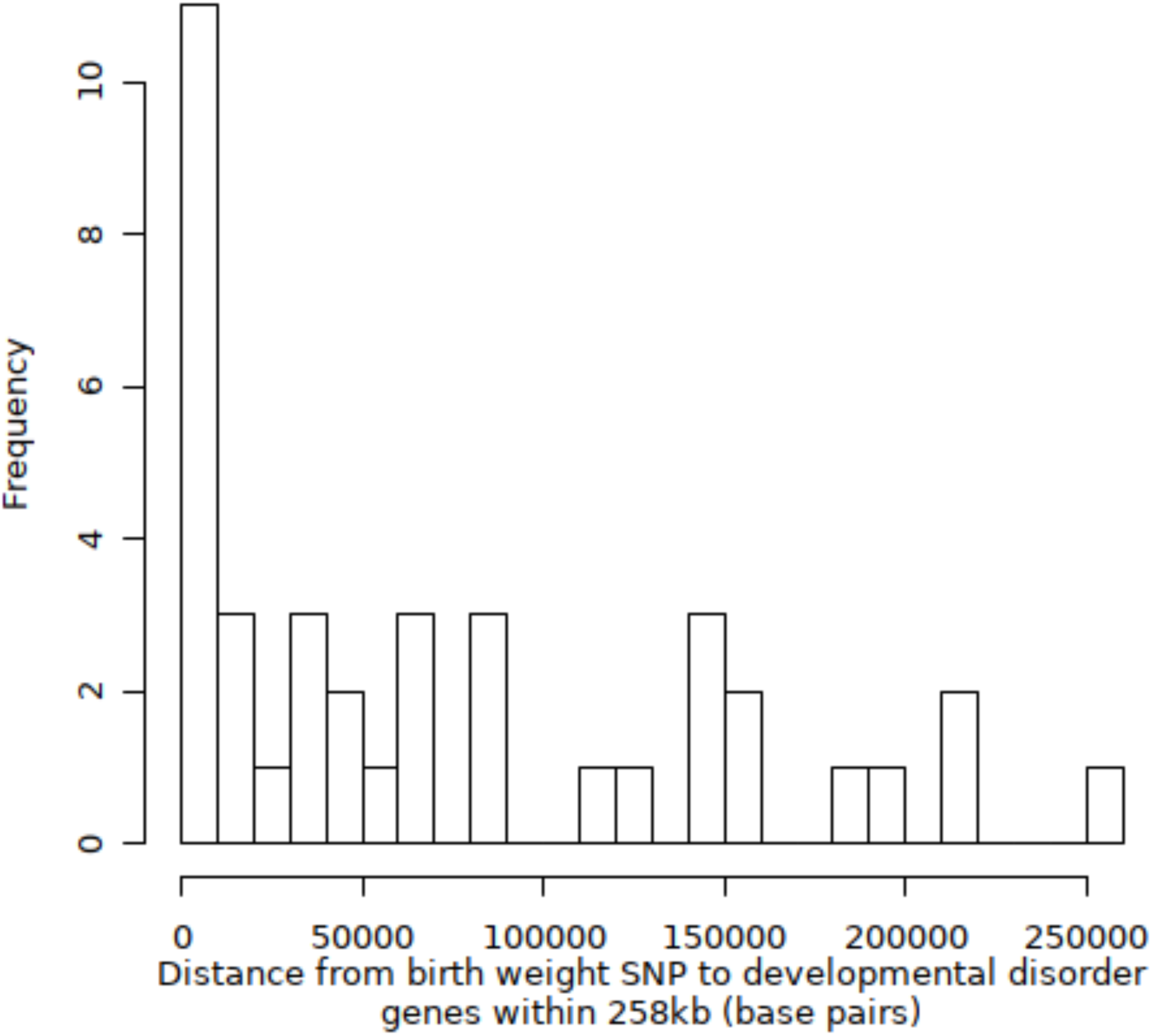
Histogram of the distance from birth weight SNPs to developmental disorder genes within 258kb of the SNP

Examples of developmental disorder genes whose associated syndromes include low or high birth weight, and which are nearby but not the nearest gene to the birth weight SNP, are *CDKN1C* and *GNAS. CDKN1C* is implicated in syndromes associated with intrauterine growth restriction (IUGR) (IMAGE syndrome) [25] and over-growth (Beckwith-Wiedemann syndrome) [26]; one of the birth weight SNPs is located 47,146bp from this gene within an intron of *KCNQ1*, which is not linked to developmental disorders. Beckwith-Wiedemann syndrome can be caused by disorders of methylation affecting imprinted genes within chromosome 11p15.5 containing *IGF2* and *CDKN1C*, both of which appeared in our analyses (Figure 3). *GNAS* has also been implicated in fetal growth, with mutations in the paternally inherited copy of the *GNAS* gene shown to lead to severe IUGR [27], and loss of methylation leading to increased fetal growth [28]. Rare mutations in this gene are also linked with low birth weight in the DECIPHER database [29] (https://decipher.sanger.ac.uk/gene/GNAS#overview/clinical-info), but the closest birth weight SNP to *GNAS* is 142,178bp away, within the *NPEPL1* gene (Figure 3). These findings support the hypothesis that the nearest gene to a SNP identified via GWAS may not always be the biologically relevant gene [30]. Syndromes resulting in large changes in birth weight associated with both of these developmental disorder genes also feature disorders of imprinting. Imprinted genes have previously been found to be enriched for birth weight associations [5], but so far no parent-of-origin specific associations have been identified at individual loci. Our approach highlights these genes as potential candidates for identifying imprinting effects affecting birth weight within the normal range.

**Figure 3:**
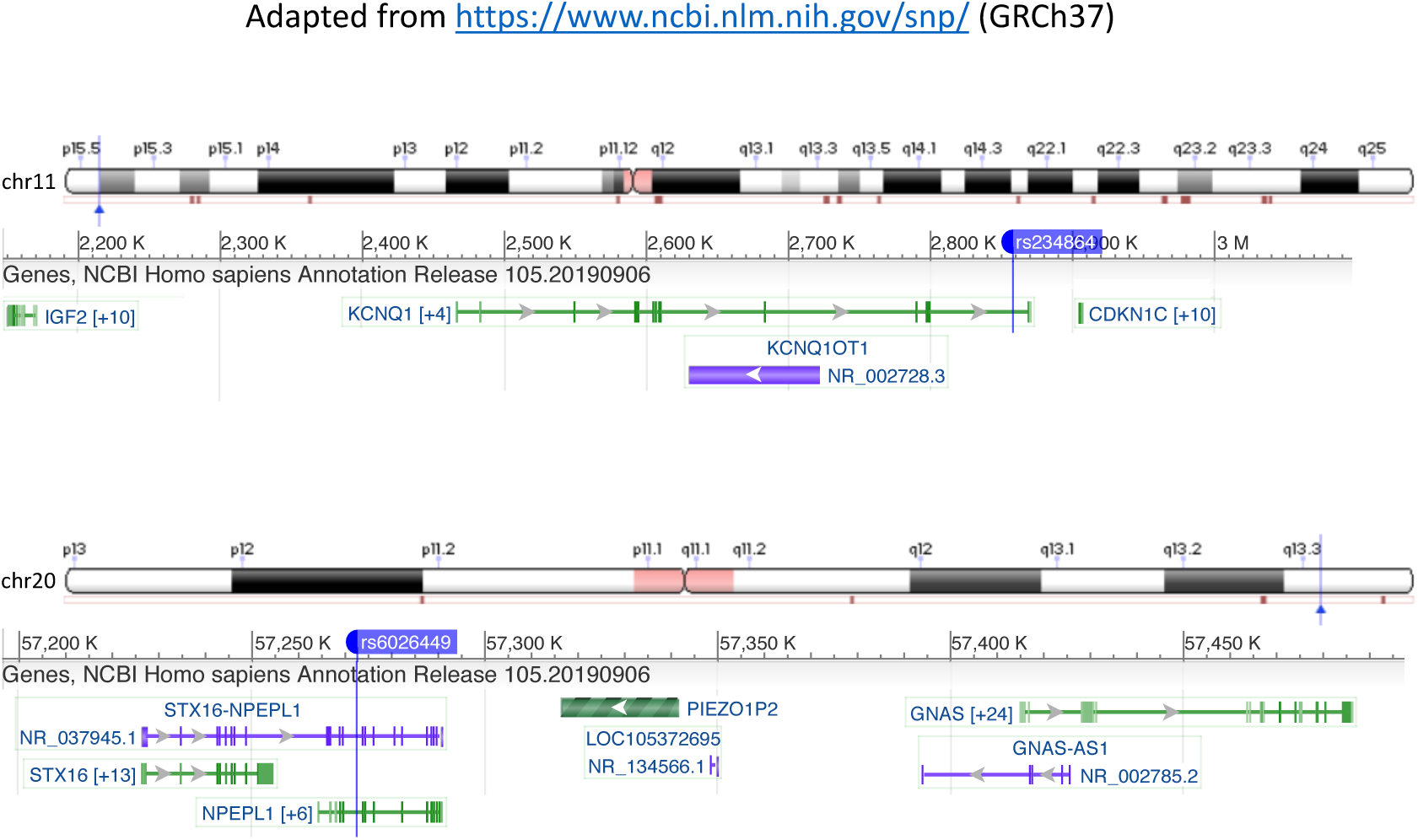
Regions surrounding the *CDKN1C* and *GNAS* genes showing the location of the genes and the nearby birth weight SNP. Colours represent different functional annotations and numbers in square brackets indicate additional transcripts.

Candidate genes highlighted by our analyses can also point towards relevant biological pathways. For example, they include three genes that are linked by IGF-1 receptor signalling (*PIK3R1, IGF1R* and *IGF2*). Two of these genes (*PIK3R1* and *IGF1R*) have one and two birth weight lead SNPs within the boundaries of the genes respectively, while the third gene (*IGF2*) is 31,481bp from the nearest birth weight lead SNP. Developmental disorders caused by variation in each of these genes are all characterised by severe effects on fetal growth. Mutations in *PIK3R* are associated with SHORT syndrome which is characterised by IUGR [31–33], and mutations causing dysregulation of *IGF1R* can also result in IUGR [34]. The *IGF2* gene is implicated in syndromes associated with fetal under-growth (Silver-Russell syndrome) or over-growth (Beckwith-Wiedemann syndrome) [35]. Furthermore, genes in the sonic hedgehog pathway have been implicated in regulation of the *IGF1R* pathway [36]. Genes from this pathway, such as *PTCH1* and *GLI2*, appear in our developmental disorder gene list, but only *PTCH1* appears in proximity to a birth weight locus in any of our analyses. Rare mutations in *PTCH1* are associated with high birth weight in the DECIPHER database and lower levels of *PTCH1* expression in preeclamptic placenta samples has been demonstrated, with strong associations between expression levels and birth weight [36]. Enrichment for SNP associations with birth weight in pathways linked to these genes has previously been demonstrated [5], but our approach highlights individual genes within the pathway which may be particularly relevant to variation in birth weight.

A pathway which was not specifically highlighted in the recent GWAS of birth weight but has come up in our analysis is the Notch signalling pathway. Alagille Syndrome, caused by rare mutations in *JAG1* and *NOTCH2*, includes failure to thrive [37] within its phenotypic spectrum. While it is not certain whether the birth weight associated SNP near *JAG1* acts primarily via fetal or maternal mechanisms, reduced expression of *JAG1* in placentas from pregnancies complicated with preeclampsia has been observed [38]. Although no association was seen between *JAG1* levels and birth weight, preeclampsia is itself associated with reduced birth weight. Other genes in the Notch pathway, *NOTCH1, NOTCH2, DLL3* and *DLL4*, were also included in our list of developmental disorder genes. Only one of these was highlighted in any of our analyses, *NOTCH1*, in the 258kb gene-window analysis, whose expression level has not previously been linked birth weight.

In the present study we have described a method for combining information from common and rare disease genetics to help prioritise candidate genes through which GWAS loci may act. We were limited by several factors. First, the list of monogenic genes we used was clinically curated as part of the Deciphering Developmental Disorders Study [18] and included any genes implicated in developmental disorders, some of which are well known to cause extremes of birth weight whilst others do not have birth weight recorded as a feature of the associated disorder. The inclusion of genes without effects on birth weight could reduce the power of the analysis to detect associations due to the inclusion of irrelevant genes. We nonetheless chose to include these genes due to the extreme rarity of many of the disorders, and thus the limited availability of detailed phenotypes, and excluding all genes without previous evidence of birth weight would similarly reduce power. Second, the list of birth weight loci also included those categorised as “Unclassified”, some of which are likely to act solely through maternal pathways. Accurate classification of these loci would also increase the power to detect enrichment, though the results of our sensitivity analysis where these loci were excluded, while less powered, were consistent with the main analysis.

In summary, we have described a novel method for testing GWAS loci for enrichment for proximity to genes implicated in monogenic disorders and demonstrated an enrichment in birth weight GWAS loci with fetal effects for proximity to genes where rare mutations are known to cause developmental disorders. This method could help prioritise candidate variants from other GWAS to help better understand the mechanisms underlying their phenotypic effect.

## Acknowledgements

The authors would like to acknowledge the use of the University of Exeter High-Performance Computing (HPC) facility in carrying out this work. This study makes use of data generated by the DECIPHER community. A full list of centres who contributed to the generation of the data is available from https://decipher.sanger.ac.uk and via email from. Funding for the project was provided by Wellcome. R.M.F. and R.N.B. are supported by Sir Henry Dale Fellowship (Wellcome Trust and Royal Society grant: WT104150). This research has been conducted using the UK Biobank Resource under application number 7036.

## Notes

### Competing Interest Statement

The authors have declared no competing interest.

